# Whole blood transcriptional signatures of age and survival identified in Long Life Family and Integrative Longevity Omics Studies

**DOI:** 10.1101/2025.07.15.664976

**Authors:** Mengze Li, Zeyuan Song, Eric Reed, Tanya T. Karagiannis, Stacy Andersen, Michael Brent, Chase Mateusiak, Sandeep Acharya, Wooseok J. Jung, Shu Liao, Mary K. Wojczynski, Mary F. Feitosa, Jeffrey R. O’Connell, May E. Montasser, Roland J. Thorpe, Konstantin Arbeev, Sofiya Milman, Albert Tai, Thomas T. Perls, Paola Sebastiani, Stefano Monti

## Abstract

Although aging is a universal event, some individuals are able to achieve extreme longevity. The Long-Life Family Study (LLFS) enrolls participants from families enriched with long-lived individuals, serves as a valuable dataset for studying ageing phenotypes and identify potential intervention targets. We analyzed the association between age at blood draw and 16,284 RNAseq-based blood transcriptomic data from 2,167 LLFS participants with ages ranging from 18 to 107, replicated the results in the Integrative Longevity Omics Study (ILO) dataset of 20,884 RNAseq-based blood transcriptomic data from 419 participants, with ages ranging from 60 to 108, and further compared our findings to a published reference aging signature.

We identified 4,227 transcripts increasing and 4,044 transcripts decreasing with age, and enrichment analysis revealed age-related upregulation of inflammatory and senescence-related pathways, and downregulation of MYC and Wnt/β-catenin targets, among others. Further, a subset of transcripts showed age associations unique to the longevity-enriched cohorts (LLFS and ILO). We also identified 314 transcripts significantly associated with mortality risk and found that pro-survival gene sets included NK cell-mediated cytotoxicity and GPCR signaling. Finally, increased transcriptomic age predicted using transcriptomic clock was strongly associated with increased mortality. In summary, this study identified robust transcriptomic signatures of aging and mortality in a longevity-enriched population, highlighting key biological pathways such as immune modulation, inflammation, and senescence.

**Author’s notes:** This manuscript has been peer-reviewed and accepted by GeroScience (Springer). This bioRxiv article reflects the published version, incorporating revisions made in response to reviewers’ comments. The main content, results, and conclusions remain unchanged from the previous version, while the Discussion section has been expanded to further address the functional annotation and clinical relevance of the identified markers.

**Copy of the acceptance letter:** The editors are pleased to inform you that your manuscript, JAAA-D-25-01946R1 entitled "Whole blood transcriptional signatures of age and survival identified in Long Life Family and Integrative Longevity Omics Studies" has been accepted for publication in GeroScience as an Original Article. The editors commend you on your outstanding contribution to the journal. Your manuscript can be published online ahead of print within approximately two weeks.

## Introduction

The biology of aging is complex and multifaceted [1] [2], with age-related changes observed across various hallmarks and omics-based profiles [1] [2] [3]. Given the growing elderly population and the increased risk for morbidity at older ages [4], significant efforts have been devoted to better understand human aging at the molecular level and to identify interventions that can maintain health [5]. Blood is a minimally invasive and reliable source of biomarkers for research and clinical diagnostics and many markers have been identified in plasma or serum [6], [7] [8]. Additionally, single-cell analyses have shown that the blood cell-type composition changes substantially during aging [9].

Aging is associated with increased risk for morbidity and mortality. However, some individuals can postpone or delay age-related diseases and extend their health span to extreme old age [10] [11], [12], [13]. Evidence is growing that these individuals harbor a combination of genetic and genomic features that either promote or reflect the biological mechanisms underlying this phenomenon. For example, we discovered unique protein signatures in centenarians [14], and distinctive metabolic profiles in long lived individuals [8]. Whole-genome sequencing analysis of semi-supercentenarians and supercentenarians has uncovered unique somatic genetic mutations in these individuals [15]. These findings underscore the value of studying long-lived individuals to uncover protective mechanisms to achieve longevity. As a critical omics layer, transcriptomic profiling holds significant potential to provide valuable insights into the biological mechanisms of aging and to identify potential therapeutic targets.

A previous study of the blood transcriptome of 14,983 individuals of European ancestry by Peters et al., identified 1,497 genes differentially expressed with chronological age, and found these genes to be enriched for the presence of potentially functional CpG-methylation sites in enhancer and insulator regions [7]. A second study of blood specimens of 3,388 adult individuals showed the ability of the transcriptome to identify molecular sub-types distinguished by their sex, age, and immune and metabolic status [16]. In all of these studies, the analysis was conducted on age ranges typical of the general population, thus excluding extreme old age individuals. Expanding these analyses to include long-lived subjects could offer valuable insights into the protective mechanisms underlying healthy aging and extreme longevity. The Long Life Family Study (LLFS) has collected whole blood transcriptomic data from individuals in families enriched with long-lived members, offering a unique opportunity to study the transcriptional changes associated with extreme old age.

An emerging and related area of research is the evaluation of biological age as distinct from chronological age, recognizing that the rate of aging varies among individuals and that chronological age is not always a reliable indicator of health status. To address this question, omics-based aging clocks have been developed. Further exploration of the potential of transcriptomic clocks to measure various aspects of aging and extreme longevity in humans could provide valuable insights into the biological mechanisms associated with and/or contributing to these processes.

In this study, we analyzed the whole blood transcriptomic profile of 2,167 individuals from longevity-enriched families, with the aim of identifying new gene markers and leading biological pathways regarding human aging, extreme longevity, and mortality risk. Additionally, a transcriptomic aging clock was constructed using elastic net and was bias-corrected [17], and the predicted age acceleration was analyzed in relation to mortality risk.

## Methods

### Study participants

The Long Life Family Study (LLFS) enrolled families with evidence of exceptional longevity across the United States and Denmark between 2006 and 2009. Study participants are followed longitudinally until death. We described study design, genetic, genomic and phenotypic data previously [18]. The Integrative Longevity Omics (ILO) study enrolled ∼1400 centenarians, their biological offspring, and spouses of the offspring between 2019 and 2024 from North America. Study participants completed a comprehensive assessment of physical and cognitive functions, and answered questionnaires on medical history and medication use, which are harmonized to LLFS. LLFS was approved by the Washington University in St. Louis IRB. ILO was approved by Albert Einstein College of Medicine IRB. All participants provided informed consent.

### RNA-seq data generation and QC

The RNA-seq data in the LLFS was generated from March 2021 to November 2023 at Washington University in St. Louis. Details are provided in [19]. Briefly, total RNA was extracted from PAXgene™ Blood RNA tubes using the Qiagen PreAnalytiX PAXgene Blood miRNA Kit (Qiagen, Valencia, CA). The Qiagen QIAcube extraction robot performed the extraction according to the company’s protocol. The RNA-Seq data were processed with the nf-core/rnaseq pipeline version 3.3 using STAR/RSEM [20].

Samples with high intergenic read percentage (> 8%), suspicious sex chromosome gene expression, or without sufficient genotype or phenotype information were excluded. Transcripts with less than 10 counts per million in at least 3% of samples across the original 4189-sample series were excluded, yielding a final set of 16,284 transcripts from 2,167 subjects for the downstream analysis.

The RNA-seq profiles of 434 ILO participant blood samples were generated from December 2023 to February 2024 in 10 batches, prepared on separate days, at the Tufts University Core Facility Genomics Core in Boston, MA. Total RNA was extracted from PAXgene™ Blood RNA tubes using the QIASymphony PAXgene Blood RNA Kit (Qiagen, Valencia, CA). The Qiagen QIASymphony SP/AS nucleic acid isolation robotic system performed the extraction according to the company’s protocol. Quality of the isolated RNA samples was checked by measuring their RNA quality numbers and concentrations using the Agilent Fragment Analyzer (Agilent, Santa Clara, CA). RNA-seq libraries were prepared using the Illumina Stranded Total RNA with RiboZero Plus preparation kit (Illumina, San Diego, CA). RNA-seq libraries were sequenced using the Illumina NovaSeq X Plus (Illumina, San Diego, CA), generating 150-base paired-end reads. Raw RNA-seq reads were processed using the nf-core/rnaseq pipeline (version 3.14) [20]. Main steps included adapter trimming, quality control, and alignment steps. Nextera transposase adapter sequences were removed using TrimGalore (version 0.6.7) [21] [22]. Trimmed reads were aligned to the human reference genome, GRCh38, using STAR version 2.7.9a [23], followed by gene expression quantification using Salmon version 1.10.1 [24]. In addition, the genomic origins of all reads, which could be one of exonic, intronic, or intergenic, were quantified using Qualimap version 2.3 [25]. Following read processing, low-quality samples were flagged based on various quality control metrics, including total reads, total aligned reads, total reads counted in genes, percentage of total reads that were aligned, percentage of aligned reads that were counted, percentage of intergenic reads, and the total number of genes with non-zero counts. For each metric, samples were flagged based on interquartile range outlier detection of all samples, and samples that flagged as outliers in multiple metrics were excluded from further analysis. As a result, 419 of the 434 profiles were retained. Following sample filtering, transcripts were excluded if they were poorly quantified in any of the 10 batches, defined as having zero counts in at least 20% of samples within at least one batch. As a result, 20,884 of the 57,890 transcripts were retained.

For both LLFS and ILO transcriptomic data, raw transcript counts were normalized using the R package *DESeq2* [26] and log2-transformed.

### Genotype data

To control for population structure and familial relatedness among the study participants, we calculated genome-wide principal components (PCs) and genetic relationship matrix (GRM) from the LLFS whole genome sequencing data released in December 2021 using the pipeline described in [27]. The genetic data were described in [28].

### Transcriptomic signature of chronological age

In LLFS data, we applied a linear mixed-effects model to regress each transcript’s expression level against age at blood draw using the *fitNullModel* function in the R package *GENESIS* [29]. The model included a random intercept and specified the variance-covariance structure of the random effects using the GRM to account for participant relatedness. The initial dataset comprised 2,167 participants; after excluding individuals with incomplete covariate data, we retained 2,101 participants from 183 families for the final analysis (Table 1). Each gene-level model incorporated covariate adjustments for sex, education level, recruitment center, RNA-seq instrument batch, percentage of intergenic reads, and the first ten genotype-derived principal components.

**Table 1.**
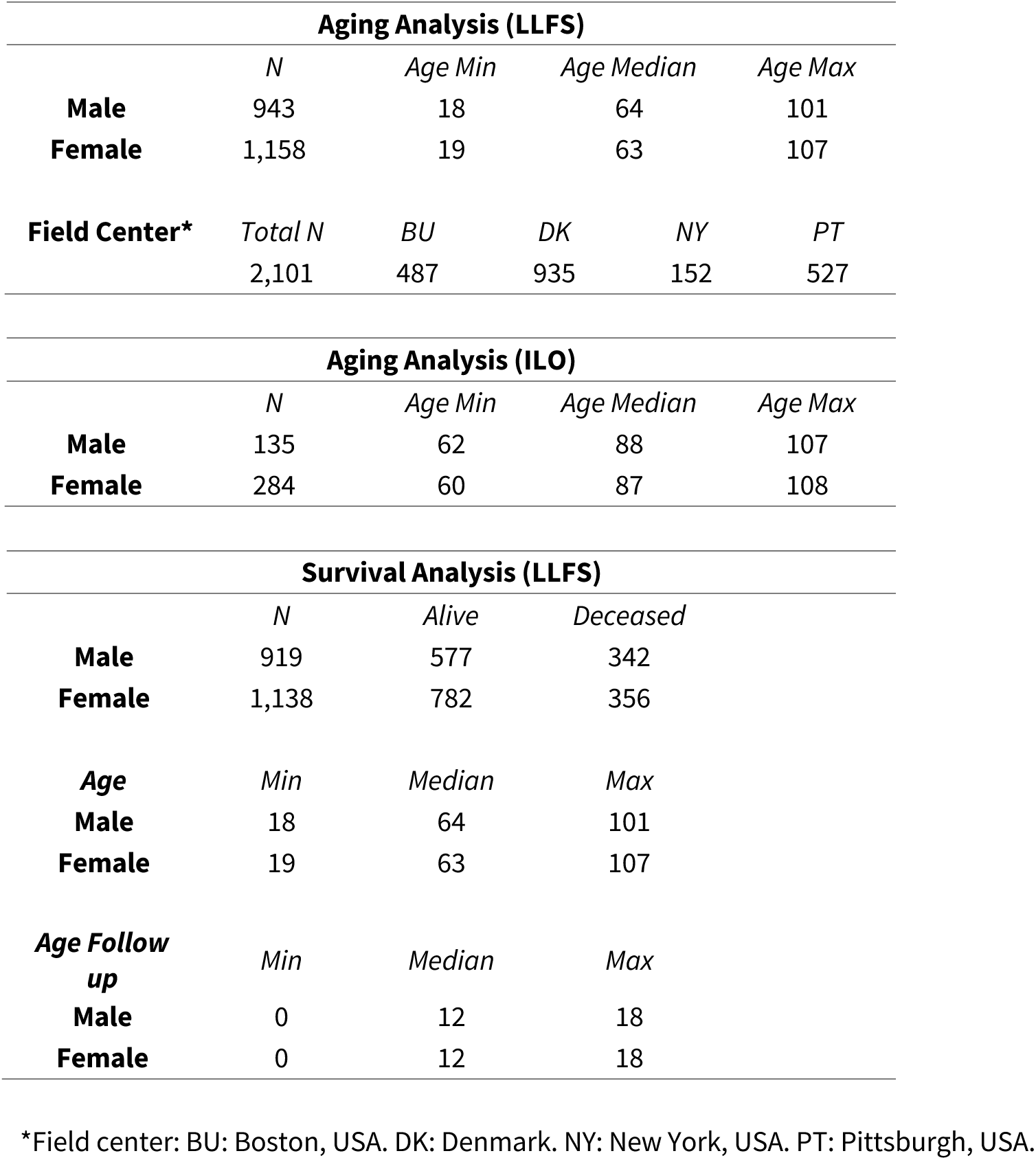
Characteristics of the study participants.

| Aging Analysis (LLFS) |  |  |  |  |  |
| --- | --- | --- | --- | --- | --- |
|  | *N* | *Age Min* | *Age Median* | *Age Max* |  |
| Male | 943 | 18 | 64 | 101 |  |
| Female | 1,158 | 19 | 63 | 107 |  |
| Field Center* | *Total N* | *BU* | *DK* | *NY* | *PT* |
|  | 2,101 | 487 | 935 | 152 | 527 |
| Aging Analysis (ILO) |  |  |  |  |  |
|  | *N* | *Age Min* | *Age Median* | *Age Max* |  |
| Male | 135 | 62 | 88 | 107 |  |
| Female | 284 | 60 | 87 | 108 |  |
| Survival Analysis (LLFS) |  |  |  |  |  |
|  | *N* | *Alive* | *Deceased* |  |  |
| Male | 919 | 577 | 342 |  |  |
| Female | 1,138 | 782 | 356 |  |  |
| Age | *Min* | *Median* | *Max* |  |  |
| Male | 18 | 64 | 101 |  |  |
| Female | 19 | 63 | 107 |  |  |
| Age Follow up | *Min* | *Median* | *Max* |  |  |
| Male | 0 | 12 | 18 |  |  |
| Female | 0 | 12 | 18 |  |  |

\*Field center: BU: Boston, USA. DK: Denmark. NY: New York, USA. PT: Pittsburgh, USA.

1,786 participants had medication information available, indicating whether they were taking medications for hypertension, heart disease, high cholesterol, and diabetes (Figure S1 A). To assess the impact of medication on gene expression levels, we repeated the analysis, including these four medication groups as additional covariates to assess their effects on the association between gene expression and age. The results showed no significant effect of medication on gene expression (Figure S1B, C). Therefore, medication was not included as a covariate in the subsequent analyses.

In ILO data, we applied a linear model to regress each transcript’s expression levels against age at blood draw using the *glm()* base R function. Each gene-level model incorporated covariate adjustments for sex, education level, profile preparation date batch, and percentage of intergenic reads. For each gene, profiles with outlier expression levels were excluded from modeling, which was defined as exceeding four standard deviations from the 20% trimmed mean expression of all profiles.

For both analyses of LLFS and ILO transcriptomic data, we corrected nominal p-values for multiple testing using the Benjamini-Hochberg false discovery rate (FDR) procedure (q-values).

### Replication of the transcriptomic signature of chronological age

We compared our results with the transcriptomic signature of age reported by Peters et.al. [7] and the dataset in the ILO study. Genes were then categorized according to whether they increased or decreased with age, and Fisher’s exact tests were performed separately for the two groups to evaluate the significance of the overlap between the marker genes in the two datasets. The background for both tests was defined as the intersection of genes available in both studies. When comparing our study to Peters et.al., this overlap included 9,463 genes; for the comparison with ILO, the overlap was 14,728 genes. The null hypothesis of Fisher’s exact test is that a gene’s association with age in one study is independent of its association in the other study. Therefore, a significant deviation from this null hypothesis would indicate a successful replication of age-associated gene markers between the two studies.

### Transcriptomic signature of mortality risk

We conducted a survival analysis to identify gene transcripts that predict mortality risk. This analysis included 2,057 participants, 698 of whom were deceased (Table 1). A Cox-proportional hazards (Cox-PH) model was fitted for each gene transcript data using the *coxph* function from the R package *survival* [30]. Pedigree ID was included as the cluster term to account for family membership. In this analysis, we used time to death as the outcome of interest, and batch-corrected and standardized gene expression data as the predictor. We adjusted the model by age at enrollment, sex, education level, enrollment center, and percentage of intergenic reads in the RNA-seq experiment. To obtain the batch-corrected gene expression values, we first fitted a linear regression model for each gene, using the batch indicator as a predictor. The residuals from this model were then extracted and standardized to have a mean of 0 and a standard deviation of 1.

### Gene set enrichment analysis

We mapped Ensembl IDs of the mRNAs to gene symbols using biomaRt [31] and retrieved gene set collections from MSigDB [32], including HALLMARK, KEGG, and REACTOME. Gene set enrichment analysis was performed with hypeR [33] using a hypergeometric test. Gene sets with q ≤ 0.05 after FDR correction were considered significant and retained. We applied fgsea [34] to perform the rank-based Kolmogorov-Smirnov (KS) test, where the gene sets were ranked using their Z-score. Additionally, to specifically assess cellular senescence, we tested enrichment of the SenMayo gene set [35] with the KS test using fgsea. There were 125 genes included in the SenMayo gene set, and 67 of them passed the QC filter in the RNA-seq preprocessing and used for enrichment analysis.

### Co-expression analysis

We used Weighted Gene Co-expression Network Analysis (WGCNA) [36] to detect gene co-expression modules among the 2,550 genes most largely affected by age (absolute Z-score > 6, q ≤ 0.01). We first obtained residuals by regressing each transcript on age, sex, education level, recruitment center, RNA-seq batch number, and percentage of intergenic reads in the RNA-seq experiment. We then performed module detection using these residuals. The clustering method was selected to be “average”, and the correlation type was set to be “signed”. We defined the hub gene in each module as the gene that has the highest correlation with the module’s eigengene.

### Construction of a transcriptomic aging clock

We generated an aging clock using standardized (mean = 0, SD = 1) RNA expression levels, with chronological age as the response variable. We trained the model on data from 2,167 participants, using the elastic net method implemented in the R package *glmnet* [37]. We used cross-validation to determine optimal α and λ values, with alpha values ranging between 0 and 1 in increments of 0.05. The model yielding the lowest RMSE was chosen, corresponding to α = 0.05 and λ = 2.215. The importance of each of the genes included in the model was estimated using the permutation method, as implemented in the R package *vip* [38]. Predicted ages by the model showed a systematic bias, with younger individuals being overestimated in predicted age and older individuals being underestimated. To address this bias, we employed the correction proposed by Cole et al. [17], [39].

Survival analysis based on a Cox-PH model was performed using delta-age, defined as the difference between corrected predicted age and chronological age, as the regression variable, with a positive (negative) value predicting the subject to be biologically older (younger). The same set of covariates used in the survival signature analysis was included in the model.

## Results

### Participants’ characteristics

Pre-processing and quality-control procedures yielded robust quantification of 16,284 transcripts across 2,167 individuals from 184 families in the LLFS cohort, with participant ages ranging from 18 to 107 years (Table 1). The ILO cohort comprised 419 individuals with ages ranging from 60 to 108 years (Table 1).

### Chronological age impacts the expression of multiple genes

In the LLFS cohort, we identified 4,227 age-associated mRNA transcripts mapping to 3,777 genes whose expression increased with age, and 4,044 transcripts mapping to 3,710 genes whose expression decreased with age (1% FDR, corrected using Benjamini-Hochberg false discovery rate correction, Figure 1). The top 20 genes associated with age are shown in Table 2 and the results for all genes are available in Table S1.

**Figure 1.**
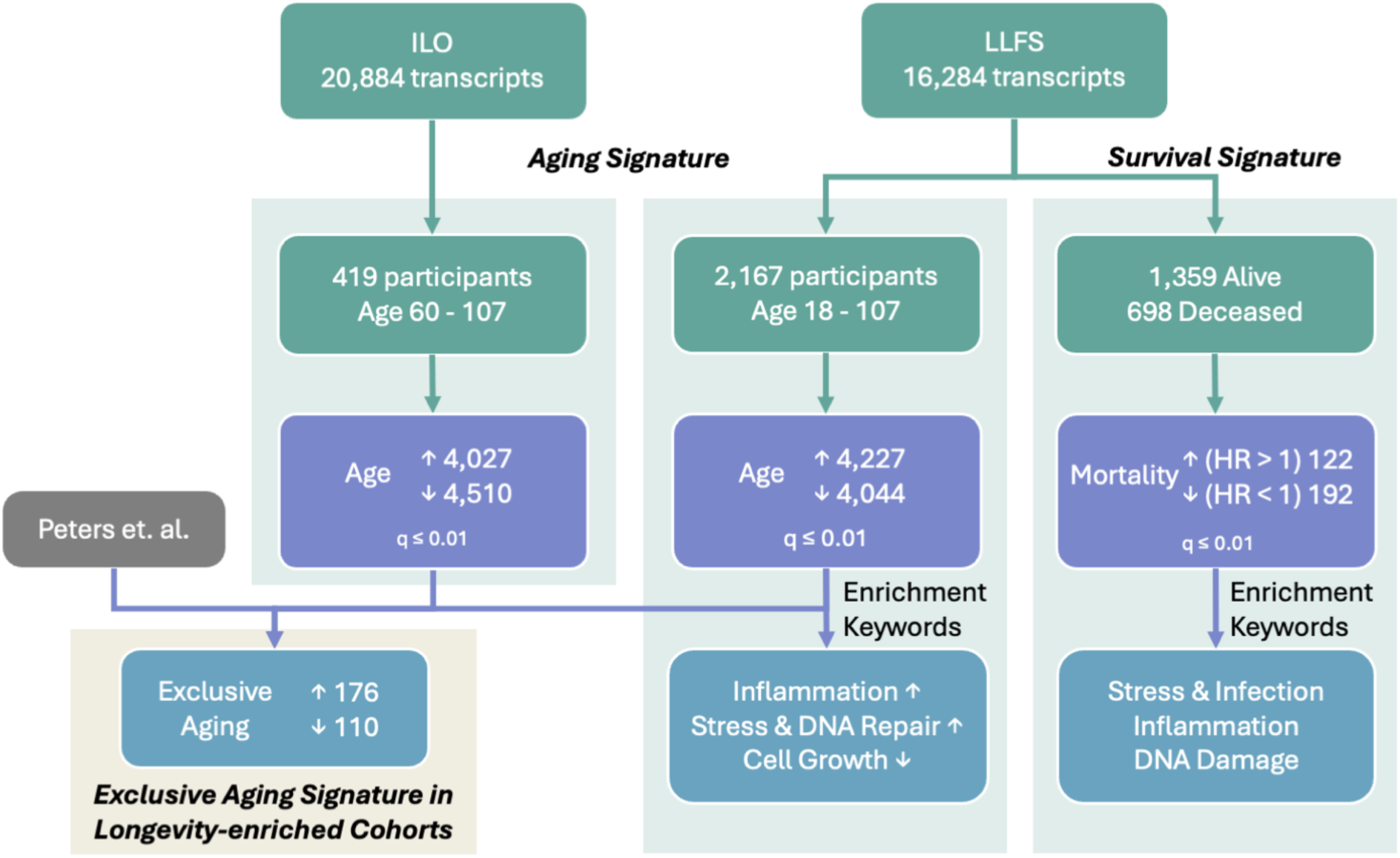
Summary of sample size and identified gene markers.

**Table 2.** Top 20 age associated genes, ranked by p-value.

| *Ensembl_ID* | *symbol* | *Age_Est* | *Age_SE* | *Age_Z* | *Age_p* | *Age_q* |
| --- | --- | --- | --- | --- | --- | --- |
| ENSG00000174807.4 | CD248 | -0.0476 | 0.00165 | -28.9 | 2.95E-183 | 4.81E-179 |
| ENSG00000173114.13 | LRRN3 | -0.0393 | 0.0016 | -24.5 | 5.92E-133 | 4.82E-129 |
| ENSG00000135318.12 | NT5E | -0.027 | 0.00128 | -21.1 | 6.53E-99 | 3.55E-95 |
| ENSG00000145911.6 | N4BP3 | -0.0219 | 0.00105 | -20.8 | 2.57E-96 | 1.05E-92 |
| ENSG00000183691.6 | NOG | -0.0347 | 0.00178 | -19.5 | 9.01E-85 | 2.93E-81 |
| ENSG00000112486.17 | CCR6 | -0.0202 | 0.00107 | -18.9 | 7.48E-80 | 2.03E-76 |
| ENSG00000081059.20 | TCF7 | -0.0146 | 0.000783 | -18.7 | 7.83E-78 | 1.82E-74 |
| ENSG00000099204.20 | ABLIM1 | -0.0142 | 0.000791 | -18 | 1.71E-72 | 3.47E-69 |
| ENSG00000184613.11 | NELL2 | -0.0207 | 0.00116 | -17.8 | 7.48E-71 | 1.35E-67 |
| ENSG00000167106.12 | EEIG1 | -0.015 | 0.000843 | -17.7 | 2.03E-70 | 3.31E-67 |
| ENSG00000144290.17 | SLC4A10 | -0.0397 | 0.00225 | -17.6 | 1.21E-69 | 1.78E-66 |
| ENSG00000232021.7 | LEF1-AS1 | -0.0261 | 0.00148 | -17.6 | 1.66E-69 | 2.25E-66 |
| ENSG00000169554.22 | ZEB2 | 0.0087 | 0.000499 | 17.4 | 5.68E-68 | 7.12E-65 |
| ENSG00000139193.4 | CD27 | -0.0146 | 0.000853 | -17.1 | 1.73E-65 | 2.01E-62 |
| ENSG00000136997.21 | MYC | -0.012 | 0.000704 | -17.1 | 3.02E-65 | 3.28E-62 |
| ENSG00000215788.11 | TNFRSF25 | -0.0139 | 0.000821 | -17 | 1.70E-64 | 1.73E-61 |
| ENSG00000228956.9 | SATB1-AS1 | -0.0157 | 0.000929 | -16.9 | 4.96E-64 | 4.75E-61 |
| ENSG00000089012.14 | SIRPG | -0.0143 | 0.00085 | -16.9 | 8.61E-64 | 7.79E-61 |
| ENSG00000178562.18 | CD28 | -0.0141 | 0.000842 | -16.8 | 3.19E-63 | 2.74E-60 |
| ENSG00000126353.3 | CCR7 | -0.019 | 0.00114 | -16.6 | 5.56E-62 | 4.52E-59 |

The expression level changes with age of several notable example genes are visualized in Figure 2A, and the volcano plot illustrating all effect sizes and significance of the analyzed genes is shown in Figure 2B. These include genes decreasing with age such as CD248 (Est = -0.0476, q = 4.81e-179), implicated in extracellular matrix binding activity and cell migration; 5’-Nucleotidase Ecto (NT5E, Est = -0.027, q = 3.55e-95), pivotal in lymphocyte differentiation and the calcification process in joints and arteries; and the MYC proto-oncogene (MYC, Est = -0.012, q = 3.28e-62), acting as a proto-oncogene and contributing to cell cycle progression and apoptosis [40].

**Figure 2.**
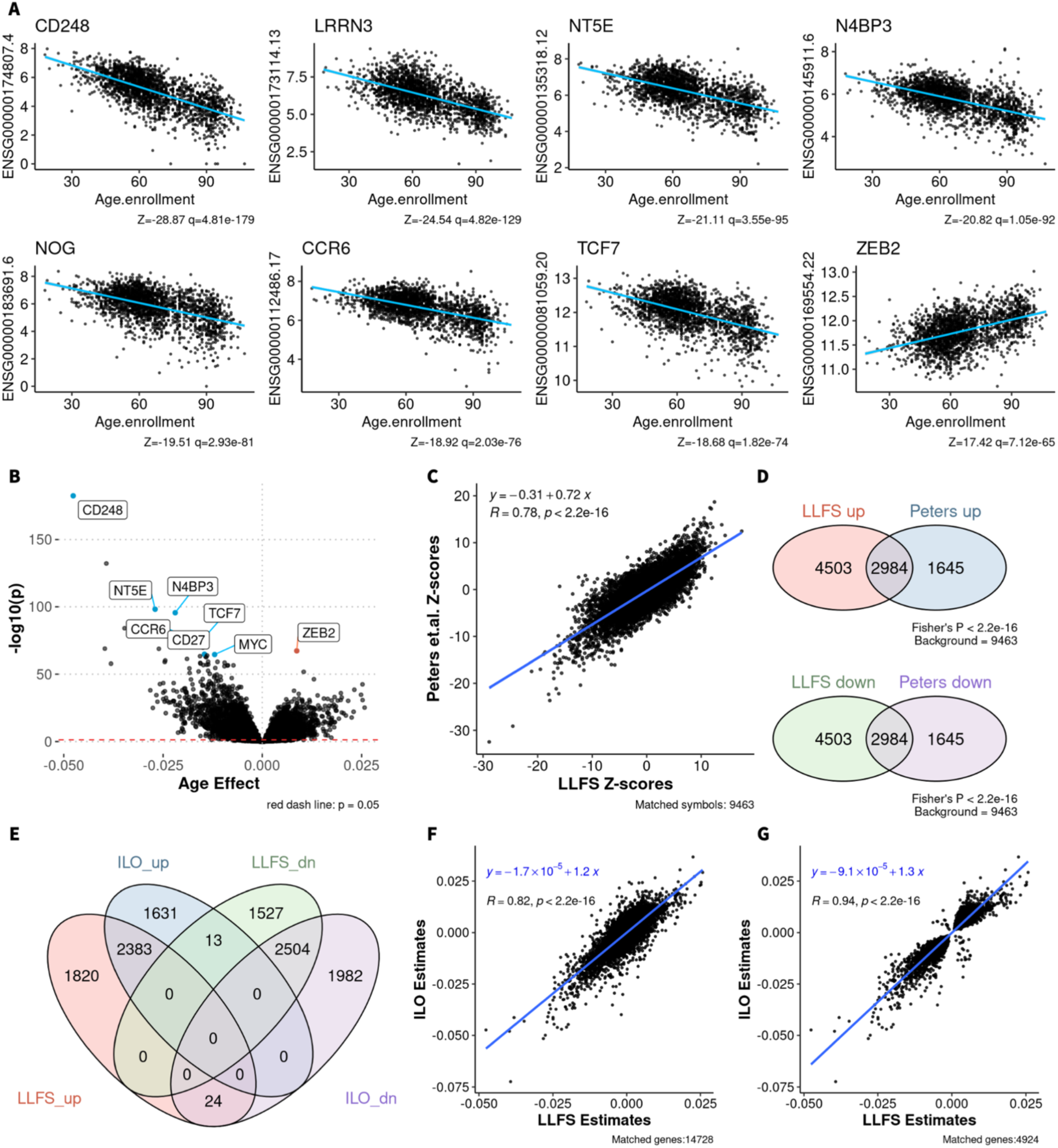
Identified age gene markers and replication. A. Scatter plots of the expression levels of selected top aging markers. Blue line represents the fitted regression line. Z-scores and q-values (FDR correction) are shown in the caption. B. Volcano plot of the analyzed gene markers. A few top markers are labeled. C. Scatter plot of Z-scores from regression analysis of age for 9,463 shared genes across LLFS and Peters.et.al. X-axis: Z-scores from LLFS. Y-axis: Z-scores from Peters et al. Blue line represents the fitted regression line, with R denoting the Pearson correlation coefficient. D. Venn diagrams of the number of significant gene markers that increase and decrease with age in LLFS and Peters et. al. Significant markers are defined by q ≤ 0.01 (FDR correction). Fisher’s exact test was conducted separately for markers increasing (up) and decreasing (down) with age, with each background set to be 9,463, which is the shared number of the genes analyzed in both studies. E. Venn diagram of the number of significant gene markers that increase and decrease with age in LLFS and ILO. Significant markers are defined by q ≤ 0.01 (FDR correction). F. Scatter plot of estimates of the effects of all the genes analyzed in both LLFS and ILO, totaling 14,728 genes. X-axis: estimates from LLFS. Y-axis: estimates from ILO. Blue line represents the fitted regression line, with R denoting the Pearson correlation coefficient. G. Scatter plot of estimates of the effects of the genes that are significantly associated with age (q ≤ 0.01) in either LLFS or ILO, totaling 4,924 genes. X-axis: estimates from LLFS. Y-axis: estimates from ILO. Blue line represents the fitted regression line, with R denoting the Pearson correlation coefficient.

Many of the most significant gene markers that decreased with age are involved in immune response and immune cell development. Examples include the NEDD4 (neural precursor cell expressed developmentally down-regulated protein 4) binding protein 3 (N4BP3, Est = -0.0219, q = 1.05e-92), involved upstream of the innate immune response and shown to regulate antiviral innate immune signaling pathways [40] [41]; the chemokine receptor 6 (CCR6, Est = -0.0202, q = 2.03e-76), expressed in various immune cells and shown to guide cell migration during immune responses [42]; the T-Cell specific transcription factor 7 (TCF7, Est = -0.0146, q = 1.82e-74), crucial for the development of natural killer cells and innate lymphoid cells; CD27 (Est = -0.0146, q = 2.01e-62), required for T cell maintenance and B cell activation; and CD28 (Est = -0.0141, q = 2.74e-60), critical for T-cell proliferation and survival [40].

In contrast, zinc finger e-box-binding homeobox 2 (ZEB2) was among the most significantly upregulated genes with age (Est = 0.0087, q = 7.12e-65). As a transcription factor, ZEB2 plays a critical role in epithelial-to-mesenchymal transition [43], is essential for the differentiation of age-associated B cells, and has been implicated in autoimmune pathogenesis [44]. Notably, studies in mice have shown that silencing Zeb2-NAT enhances the reprogramming efficiency of aged fibroblasts, underscoring its potential as a therapeutic target for preserving stem cell pluripotency [45]. Another gene, leucine rich repeat containing G protein-coupled receptor 6 (LGR6, Est = 0.0198, q = 1.53e-34), was associated with chronic activation of the Wnt/β-catenin pathway, a process that promotes cellular senescence, and exacerbates the progression of chronic obstructive pulmonary disease (COPD) [46].

### Successful replication of genes associated with age

Comparison of the aging signature we identified in LLFS with a previously published human blood transcriptomic signature of age by Peters et al. [7] showed significant replication, both in terms of correlation of the Z-scores (R = 0.78, p < 2.2e-16, Figure 2C) and in terms of signature overlap (Fisher test p < 2.2e-16, Figure 2D), confirming the robustness of our results (Table S1). The systematic bias in Z-score values can be attributed to the difference in sample size (n = 14,983 in Peters et al. and n = 2,101 in LLFS).

We performed a similar comparison with the ILO cohort and observed a strong correlation in age-related gene expression estimates across the genes shared by the two studies (R = 0.82; Figure 2F). This correlation increased to R = 0.94 when considering only the genes that were significantly associated with age (q ≤ 0.01) in either LLFS or ILO (Figure 2G). The overlap of significant gene markers identified in the two studies is also illustrated in Figure 2E.

### Functional annotation shows multiple biological processes affected by age

All analyzed gene markers were ranked by Z-score, with those showing the greatest age-related increases placed at the top and those with the greatest decreases at the bottom. Enrichment of gene sets across multiple compendia was then assessed using KS tests applying a significance threshold of q ≤ 0.05. Figure 3A displays the top enrichment results in the HALLMARK compendium, with the full results available in Table S2.

**Figure 3.**
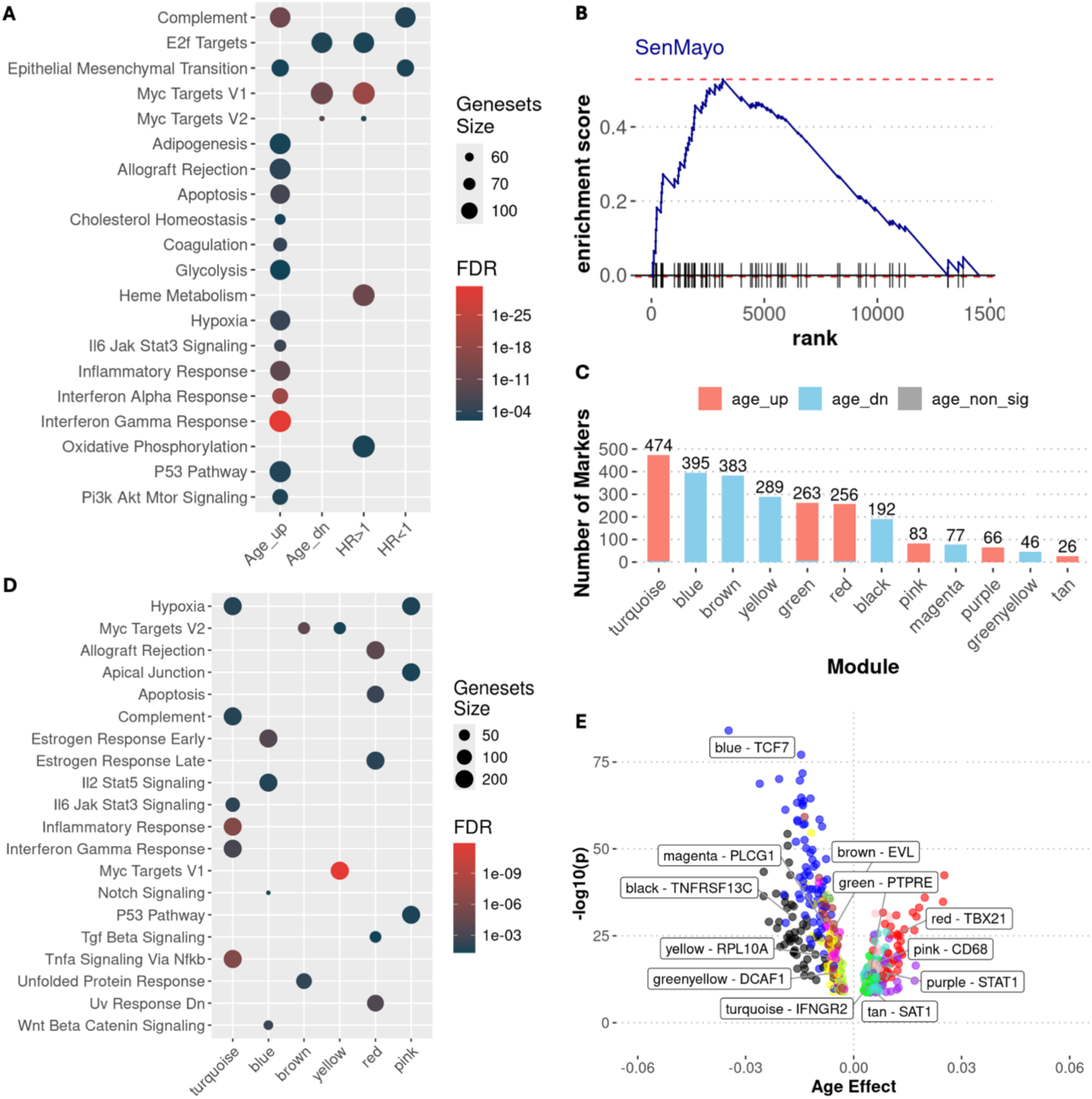
Functional annotation and co-expression module detection of age associated genes A. Gene sets from the HALLMARK compendium that significantly enriched (q ≤ 0.05) in gene markers increasing (Age_up) and decreasing (Age_dn) with age using Kolmogorov-Smirnov (KS) test. Gene markers are ranked by absolute Z score. B. Enrichment analysis of the SenMayo gene set using Kolmogorov-Smirnov (KS) test, demonstrating cellular senescence activity is enriched in markers increasing with age. Gene markers are ranked left to right based on their high-to-low age Z-scores. C to E represents the co-expression modules detected among the top aging markers (absolute Z-score > 6, q ≤ 0.01 using FDR correction) using WGCNA. C. Module sizes, with color representing the number of markers increasing and decreasing with age within the module. D. Gene sets from the HALLMARK compendium that are significantly enriched (q ≤ 0.05) in each module. Modules with no enriched gene sets are not included. E. Volcano plot of genes with correlation above 0.75 with the eigenvalue of their respective modules, with color denoting the associated module. Hub genes of each module are labeled.

The set of genes increasing with age was found to be significantly enriched in inflammation and cancer-related pathways, such as interferon gamma response (q = 3.87e-32, including PSMB10, IFI30, and VAMP5), and TNF-α signaling via NF-κB (q = 1.01e-10, including KLF10, PLEK, and CCL4). Given the reported relationship between inflammation and cell senescence [47], we further tested the enrichment of the SenMayo gene set [35] in the age signature, and showed a significant association of the gene set with markers that increase with age (enrichment score = 0.528, p = 4.39e-7, Figure 3B).

The set of genes decreasing with age was enriched in pathways related to cell growth, such as MYC targets (q = 1.83e-12), E2F targets (q = 0.0038), and Wnt/β-catenin signaling (q = 0.0129).

WGCNA-based module detection [36] of the 2,550 aging markers with absolute Z-score greater than 6 identified 12 modules of highly correlated genes, each containing between 26 and 474 genes, and with the majority of the genes in each module changing with age in the same direction (Figure 3C). The modules were further annotated by their hub genes, defined as the most ‘central’ genes within each module as measured by their correlation with the module’s first eigen value, and by gene set over-representation enrichment analysis (Figure 3D, E). This dimensionality reduction approach further highlighted dominant patterns in the aging signature.

For instance, the turquoise module, which is the largest module and primarily consists of genes that increase with age, was found to be enriched in immune-, inflammation-, and stress-related pathways, such as inflammatory response, interferon gamma response, and hypoxia (Figure 3D), indicating stress and activation of the immune system. The hub gene, interferon gamma receptor 2 (IFNGR2), is representative of the interferon pathway.

Conversely, the blue, brown, and yellow modules, which are primarily composed of genes that decrease with age, were found to be enriched in MYC targets as well as IL-2 and STAT5 signaling pathways. The hub genes for these modules were T-cell specific factor 7 (TCF7), Ena/Vasodilator-stimulated phosphoprotein-like gene (EVL), which is associated with cytoskeleton remodeling and cell migration, and Ribosomal Protein L10A (RPL10A), which encodes a ribosomal protein [40]. These findings are consistent with increased inflammation and reduced protein synthesis and cell proliferation in older individuals (Figure 3D; Table 3). Interestingly, the estrogen response early pathway was enriched in the blue module, which comprises genes that significantly decrease with age. In contrast, the estrogen response late pathway was enriched in the red module, which primarily includes genes that increase with age.

**Table 3.**
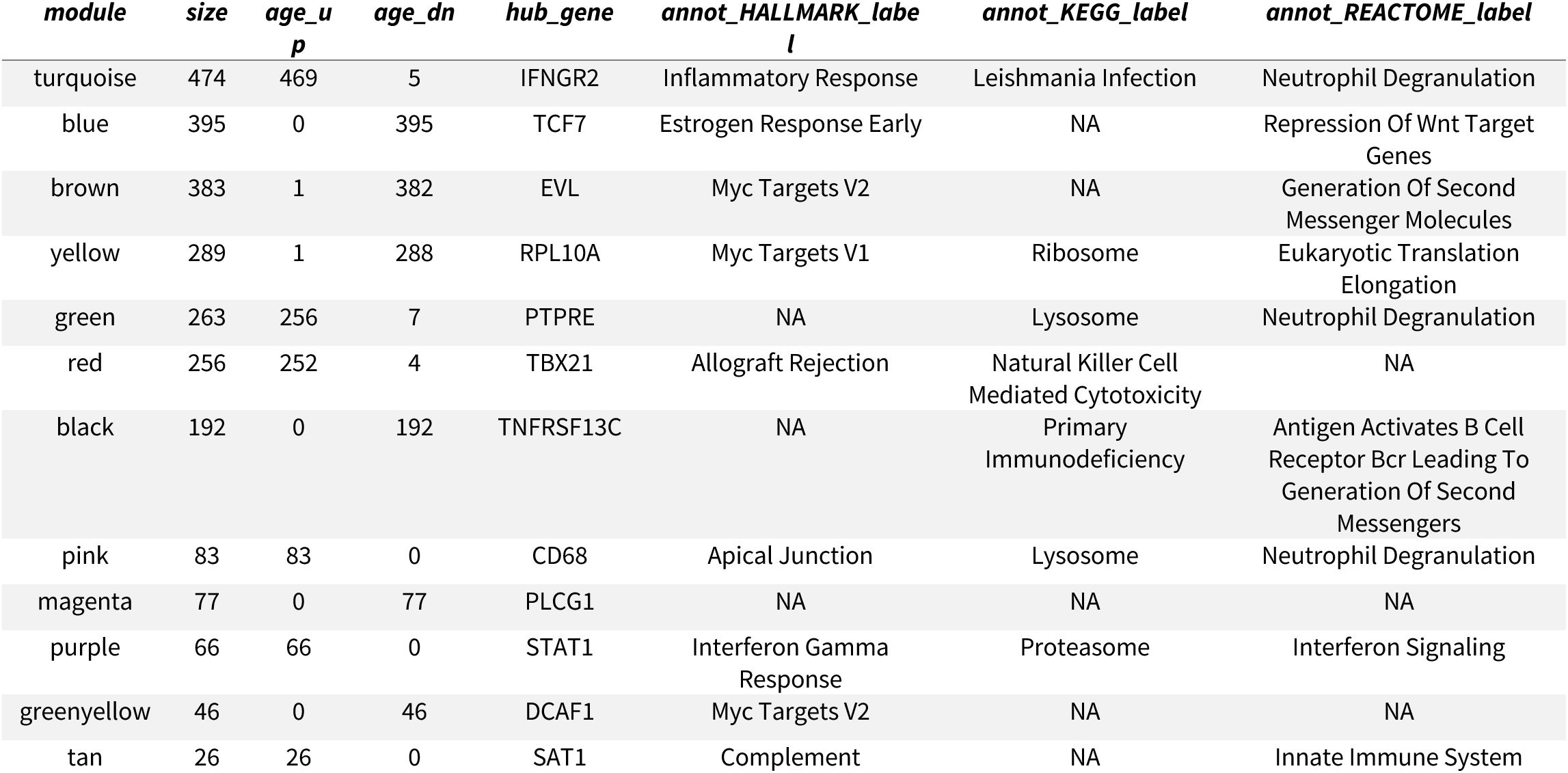
Table of modules detected from 2,550 significant transcriptomic markers (absolute Z-score > 6, q ≤ 0.01) using WGCNA, annotated by the top enriched gene sets from each compendium (HALLMARK, KEGG, and REACTOME). Only gene sets enriched with significance q < 0.05 are included. Size: the total number of markers within the module. Age_up and Age_dn: number of markers increase and decrease with age.

| *module* | *size* | *age_up*<br>*p* | *age_dn* | *hub_gene* | *annot_HALLMARK_label* | *annot_KEGG_label* | *annot_REACTOME_label* |
| --- | --- | --- | --- | --- | --- | --- | --- |
| turquoise | 474 | 469 | 5 | IFNGR2 | Inflammatory Response | Leishmania Infection | Neutrophil Degranulation |
| blue | 395 | 0 | 395 | TCF7 | Estrogen Response Early | NA | Repression Of Wnt Target Genes |
| brown | 383 | 1 | 382 | EVL | Myc Targets V2 | NA | Generation Of Second Messenger Molecules |
| yellow | 289 | 1 | 288 | RPL10A | Myc Targets V1 | Ribosome | Eukaryotic Translation Elongation |
| green | 263 | 256 | 7 | PTPRE | NA | Lysosome | Neutrophil Degranulation |
| red | 256 | 252 | 4 | TBX21 | Allograft Rejection | Natural Killer Cell Mediated Cytotoxicity | NA |
| black | 192 | 0 | 192 | TNFRSF13C | NA | Primary Immunodeficiency | Antigen Activates B Cell Receptor Bcr Leading To Generation Of Second Messengers |
| pink | 83 | 83 | 0 | CD68 | Apical Junction | Lysosome | Neutrophil Degranulation |
| magenta | 77 | 0 | 77 | PLCG1 | NA | NA | NA |
| purple | 66 | 66 | 0 | STAT1 | Interferon Gamma Response | Proteasome | Interferon Signaling |
| greenyellow | 46 | 0 | 46 | DCAF1 | Myc Targets V2 | NA | NA |
| tan | 26 | 26 | 0 | SAT1 | Complement | NA | Innate Immune System |

### Several blood transcripts predict mortality risk

We found 122 mRNA transcripts, encoding 122 genes, whose increased expression was associated with higher mortality risk (HR > 1), and 192 mRNA transcripts, encoding 175 genes, whose increased expression was associated with lower mortality risk (HR < 1, Figure 1). The full results are provided in Table S1.

KS-based enrichment analysis identified several gene sets significantly associated with mortality risk. Gene sets associated with increased mortality risk (HR > 1) included MYC targets (q = 8.67e-20), E2F targets (q = 2.74e-3), and heme metabolism (q = 1.35e-12). In contrast, gene sets associated with decreased mortality risk (HR < 1) included natural killer cell-mediated cytotoxicity (q = 2.47e-5) and signaling by GPCR (q = 1.88e-5). The full list of results is provided in Table S2.

### A subset of aging markers is uniquely associated with age in long-lived individuals

Since both LLFS and ILO cohorts are enriched for individuals who have achieved exceptional longevity (EL), we compared our findings with previously published aging signatures by Peters et al. [7], whose study did not include EL individuals, to identify aging markers potentially unique to this distinct population. Across the three studies, we observed an overlap of 9,358 gene markers. To define markers specific to the long-lived population, we selected genes whose expression levels were significantly associated with age (q ≤ 0.01) in both LLFS and ILO but not associated with age (p > 0.25) in the Peters et al. dataset. Based on these criteria, we identified 176 and 110 gene markers that uniquely increased and decreased with age, respectively, in LLFS and ILO. Notably, 14 of these genes were also significantly associated with mortality risk in LLFS (q ≤ 0.05), with 8 of them meeting a more stringent threshold of q ≤ 0.01 (Figure 4A, Figure S2). Z-score normalized expression levels of these genes in LLFS, stratified by age groups in ten-year intervals, are visualized as a heatmap in Figure S2, highlighting age-related expression changes.

**Figure 4.**
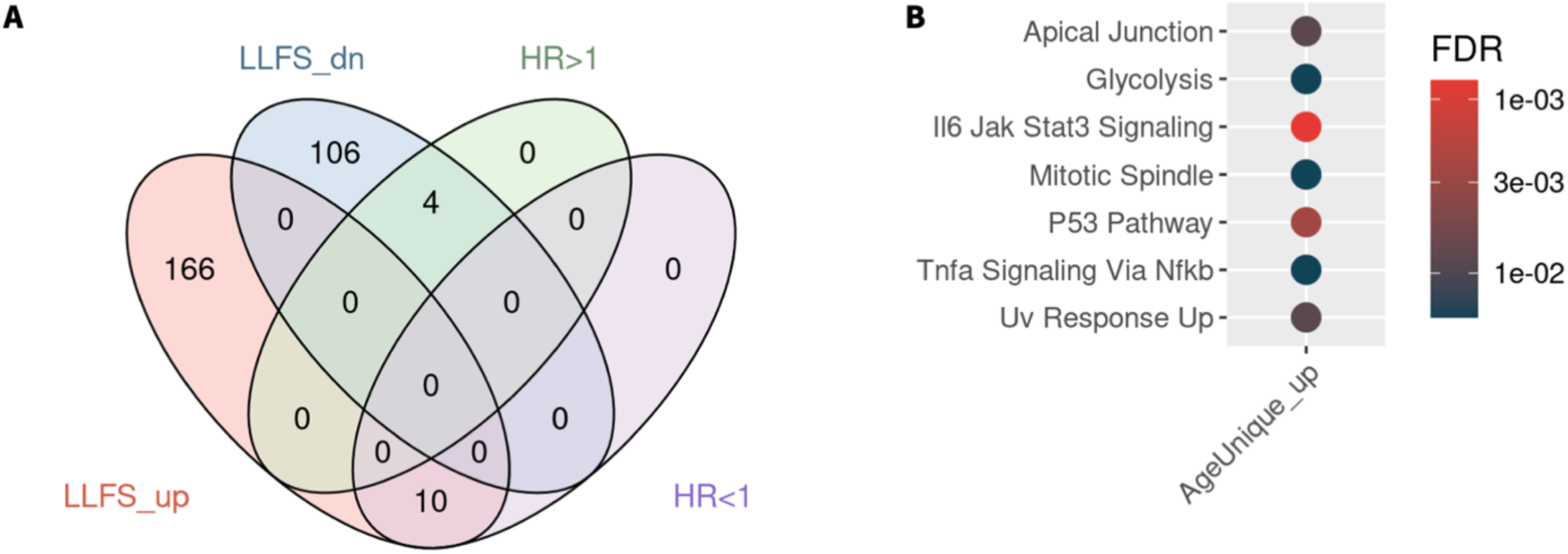
Unique markers associated with age in longevity enriched families (LLFS and ILO) A. Venn diagram illustrating the number of gene markers that are exclusively significantly associated with age in the LLFS and ILO cohorts (q ≤ 0.01), and with mortality risk (q ≤ 0.05), but not associated with age in the Peters et al. study (p > 0.25). B. Enriched gene sets among the gene markers that are exclusively associated with age in the LLFS and ILO cohorts.

Gene set enrichment analysis revealed pathways significantly associated with markers uniquely linked to age in longevity-enriched families (Figure 4B). Several pathways, such as IL6-Jak-Stat3 signaling (q = 7.8e-4) and the p53 pathway (q = 2.5e-3), overlapped with those enriched among the broader set of age-associated markers described earlier. However, additional pathways emerged in this subset, including Mitotic Spindle (q = 0.018), which was not identified in the original aging signature analysis. Other pathways, such as UV Response (q = 8.3e-3) and Glycolysis (q = 0.018), were identified in both analyses; notably, their enrichment significance remained relatively stable despite the smaller number of age-associated genes specific to longevity-enriched families.

### Age Acceleration derived from a transcriptomic clock predicts increased mortality risk

We fitted a transcriptomic clock using an elastic net model on all transcripts that passed quality control, resulting in a final model that included 1,432 transcripts with non-zero coefficients, with their distribution visualized in Figure 5A. The top genes ranked by permutation-based variable importance are shown in Figure 5B.

**Figure 5.**
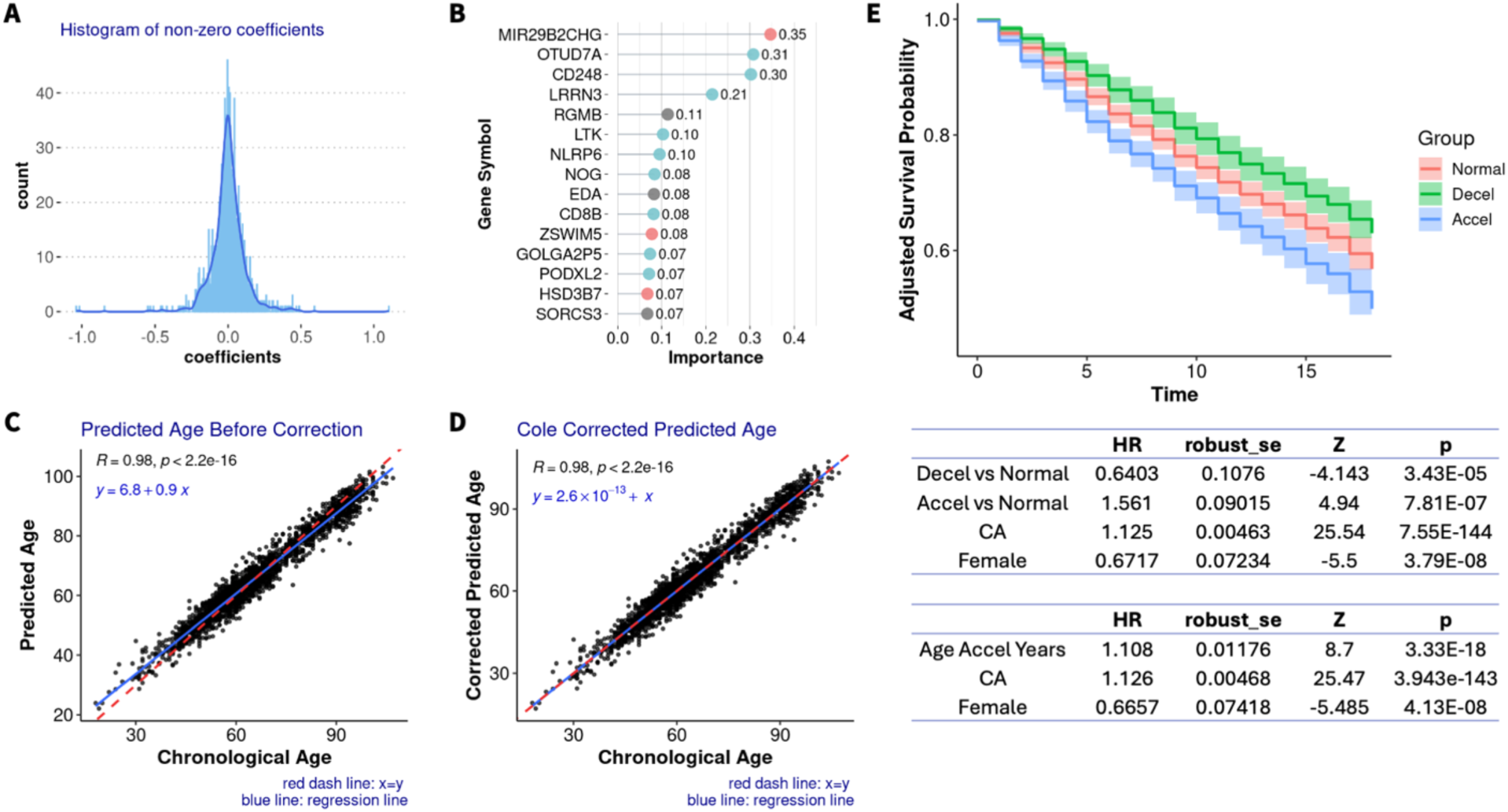
Construction of the transcriptomics aging clock and the association between age deviation (delta-age) and mortality risk. A. Histogram showing the distribution of feature coefficients. Only the 1,432 transcripts with non-zero coefficients are included. The darker blue line illustrates the density. B. Top important features in the aging clock, with importance scores derived by the permutation method. Red and green dots denote features positively (est > 0) and negatively (est < 0) associated with age (q ≤ 0.01) in the aging signature analysis, respectively. C. Comparison of predicted age versus chronological age, which shows a systematic bias in the predicted age before correction. D. Corrected predicted age (using the Cole method) versus chronological age, demonstrating the removal of the bias. E. Kaplan-Meier curve of mortality risk based on delta-age (defined by corrected predicted age minus chronological age) groups, with coefficients and significance indicated below. Normal: individuals whose corrected predicted age differs from their chronological age by no more than 3 years. Decel: individuals predicted to be over 3 years younger than their chronological age. Accel: individuals predicted to be over 3 years older than their chronological age. Age Accel Years: age acceleration as a continuous variable. CA: Chronological age. Female: Effect of female compared to male.

Upon applying the model to the dataset, a systematic bias was observed in the predicted ages (Figure 5C), which was also observed in previous studies [6], [17]. The bias was corrected using Cole’s formula [17], [39], as shown in Figure 5D. Delta-age was defined as the difference between the adjusted predicted age and chronological age, and its impact on mortality risk was evaluated both as a continuous variable and as categorical groups. Cox-PH models were applied with adjustments for relevant covariates. As a continuous variable, delta-age was significantly positively associated with mortality risk (HR = 1.108, p = 3.33e-18, Figure 5E). When recoded as a three-value categorical variable, survival analysis revealed that individuals with a negative delta-age of more than three years had a significantly lower mortality risk (HR = 0.6403, p = 3.43e-5, Figure 5E), whereas those with a positive delta-age of more than three years showed a significantly higher mortality risk (HR = 1.561, p = 7.81e-7, Figure 5E). These trends are also reflected in the age-adjusted Kaplan-Meier curve of Figure 5E.

## Discussion

In this study, we performed a large-scale transcriptomics profiling of subjects from families enriched for long-lived individuals and spanning a wide age range, between 18 and 107 years. The large sample size (n = 2,167) and the wide age range allowed us to robustly identify signatures associated with chronological age, mortality risk, and extreme longevity, and to integrate these results toward the prioritization of candidate unique aging markers.

Our analysis detected 8,271 genes exhibiting significant associations with age (q ≤ 0.01), with roughly equal numbers increasing and decreasing over the lifespan. These age-related expression changes highlight major transcriptional remodeling with age, reflecting coordinated shifts in immune function, inflammation, cellular stress responses, and cell cycle regulation. Notably, genes involved in immune cell activation and differentiation, such as CD28, CD27, TCF7, and CCR6, exhibited marked declines with age, consistent with the well-documented phenomenon of immunosenescence. Conversely, genes such as ZEB2 and LGR6 were significantly upregulated with age, suggesting increased activity in pathways related to cellular senescence, epithelial-to-mesenchymal transition, and Wnt/β-catenin signaling, which are hallmarks of aging and chronic disease progression. This finding is consistent with a previous genome-wide association study in LLFS, which also identified several genes involved in the Wnt/β-catenin signaling pathway as being associated with age-related traits [48].

Our results strongly corroborate previous findings, particularly the transcriptomic signature of aging reported by Peters et al. Despite differences in sample size and demographic composition, we observed strong replication of gene-level associations and pathway enrichments, confirming the robustness and generalizability of our aging signature. Replication across the ILO cohort further strengthens our findings and demonstrates that transcriptomic signatures of aging are largely conserved across different populations and family-based study designs.

### Enriched gene sets revealed biological processes known to play a role in aging

Functional enrichment analyses revealed that genes upregulated with age were consistently enriched in inflammatory and immune response pathways, including TNF-α signaling and interferon gamma response. This aligns with the concept of "inflammaging," a state of chronic, low-grade inflammation that contributes to age-related morbidity [1]. The enrichment of the SenMayo gene set among age-upregulated transcripts further emphasizes the role of cellular senescence in the aging process [35], as well as its link to inflammation and its established role as a hallmark of aging [1]. In contrast, genes downregulated with age were enriched for MYC and E2F targets and Wnt/β-catenin signaling, suggesting declining proliferative capacity and regenerative potential with age. Prior studies have shown that MYC also plays a role in nutrient sensing [2], and its downregulation can promote longevity and health span in mice [49]. Additionally, some of the key biological functions regulated by MYC, such as ribosome biogenesis and protein translation, are essential for cell proliferation and survival [50]. These findings highlight the interconnected roles of cell cycle regulation, nutrient sensing, and MYC signaling in the aging process.

To distill these complex changes into interpretable networks, we employed WGCNA to identify co-expression modules among highly age-associated genes. These modules revealed distinct transcriptional programs reflecting core aging processes, including immune activation, translational suppression, and cytoskeletal remodeling. For example, the turquoise module, enriched for inflammatory and interferon pathways, was anchored by the hub gene *IFNGR2*. Conversely, modules composed of downregulated genes were enriched for MYC target genes and IL-2/STAT5 signaling, with hub genes such as TCF7, EVL, and RPL10A pointing to declining immune signaling and ribosomal function.

Interestingly, our data also revealed differential enrichment of early and late estrogen response pathways, with early response genes declining and late response genes increasing with age. Though both pathways have been implicated in breast cancer biology [51] [52], their distinct age-related regulation may reflect complex hormonal changes in aging and could have implications for sex-specific disease risk.

### Mortality-associated genes offer insight into underlying biological mechanisms

Many of the identified transcripts were also predictive of mortality. Gene sets associated with increased mortality risk included MYC and E2F targets, as well as pathways related to heme metabolism, whereas protective gene sets included those involved in chemokine signaling, natural killer (NK) cell-mediated cytotoxicity, and G protein–coupled receptor (GPCR) signaling, the latter previously shown to influence cellular senescence and cell cycle control [53]. These findings suggest that a robust immune response may support the detection and clearance of abnormal or senescent cells, thereby reduce the risk of malignancy and contribute to improved survival. Conversely, our observation that elevated expression of *MYC* targets is associated with increased mortality aligns with experimental evidence that heterozygous Myc ± mice exhibit extended lifespan compared to wild type (Myc +/+) mice [54].

Elevated expression of several ribosomal proteins, including RPL32, RPL19, and RPS8, was significantly associated with increased mortality risk. These genes contribute to the activation of several pathways linked to nutrient sensing and cellular biogenesis, including the EIF2AK4 (GCN2) response to amino acid deficiency, seleno-amino acid metabolism, and cellular responses to starvation. In addition, they were enriched in the *regulation of SLITs and ROBOs expression* and *ROBO receptor signaling*, pathways previously implicated in inflammation and tissue remodeling [55].

Interestingly, these same ribosomal proteins were found to decrease in expression with chronological age. This may reflect the well-documented decline in efficient protein synthesis capacity with age [1]. However, in the context of disease or compensatory responses, dysregulation of these pathways may contribute to maladaptive cellular states, thereby increasing mortality risk. Further investigation is warranted to clarify the context-specific roles of these ribosomal genes in aging and survival.

Taken together, these findings support the potential utility of blood transcriptomic markers as prognostic indicators of biological age and mortality risk. They also underscore the importance of specific transcriptional programs – particularly those related to immune surveillance, cellular metabolism, and translational control – in mediating resilience or susceptibility to age-related decline.

### Functional insights into aging markers unique to individuals from families enriched in exceptional longevity

A particularly novel aspect of our study was the identification of a subset of aging markers that are associated with age in the LLFS and ILO cohorts, but not in the general population. These unique markers were enriched in pathways such as IL6-Jak-Stat3 signaling and p53 signaling, as well as mitotic spindle and glycolysis pathways not observed in the broader age signature. The emergence of these unique signatures in long-lived families suggests that distinct regulatory programs may underlie healthy aging and exceptional longevity.

Some of these markers may play a protective role against age-related decline. For example, regulator of G protein signaling 3 (RGS3), which increases with age only in longevity-enriched cohorts, was previously shown to confer protection against pathological changes in fibroblasts [56]. Although not yet extensively studied, it has demonstrated potential for clinical application. Members of the RGS family have been proposed as biomarkers in cancer [57], highlighting their growing relevance in clinical settings. In addition, while no studies have directly examined RGS3 in this context, it modulates the KRAS pathway [58] which is involved in cancer progression. A clinical trial for non-small cell lung cancer is currently underway (ClinicalTrials.gov ID: NCT06582771) that targets KRAS and includes RGS3 as one of its evaluation points. Taken together, a higher expression level of RGS3 seem to be beneficial as it could suppress oncogenic KRAS activity, which aligns with our hypothesis, and it has already showcased its potential application in clinical settings. Similarly, CD200, which is known to be elevated in pancreatic ductal adenocarcinoma [59], exhibited a unique age-related decrease in the longevity enriched families, suggesting a potentially beneficial regulatory role in aging within this cohort. CD200R1, the receptor for CD200, has already been identified as a promising therapeutic target in a phase 1 clinical trial [60], supporting the potential utility of CD200 in clinical applications.

In contrast, however, the functional roles of some other markers are less clear. For instance, NR1D1, a nuclear receptor with reported anti-inflammatory role through inhibition of M1 macrophage polarization via suppression of the NF-κB pathway [61], showed a decline with age. This may indicate a deleterious shift in immune regulation. However, given NR1D1’s involvement in circadian rhythm regulation, light exposure, and physical activity [62], its downregulation may also reflect environmental or behavioral changes with age, rather than intrinsic biological decline.

Similarly, PIK3R3 (est = 8.961e-3, q = 1.397e-14) and RNF114 (est = 2.871e-3, q = 3.703e-14), both of which demonstrated age-associated increases in expression, have been implicated in promoting colorectal cancer cell proliferation [63] [64], with PIK3R3 also upregulated in liver cancer cells [65]. Although these associations might initially suggest harmful effects, it is important to consider the context of these findings. Prior studies were conducted in cancer cell lines, whereas our observations are derived from whole blood transcriptomic data in non-diseased individuals. While these genes are linked to proliferative signaling in cancer, their expression in healthy aging may reflect preserved proliferative or regenerative capacity. Given that aging is typically associated with diminished cell proliferation and tissue maintenance, it is possible that increased expression of such genes may, under non-pathological conditions, support tissue repair and homeostasis, underscoring their potentially beneficial context-dependent roles.

Pathway analysis further reveals unique patterns in those families enriched with exceptional longevity. Several immune and inflammation-related pathways, including IL6-Jak-Stat3 signaling, the p53 pathway, and TNF-α signaling via NF-κB, were enriched both in this subset and in the broader aging marker analysis (Figure 3A). However, additional pathways emerged as unique to the long-lived population. Notably, the UV response up pathway, associated with external stress, was enriched among genes that uniquely increased with age. Additionally, enrichment of glycolysis and the mitotic spindle pathways, which are involved in energy metabolism and cell division, suggests heightened cellular activity and proliferative potential in individuals from families with exceptional longevity.

### Delta-age predicted using a transcriptomics aging clock is associated with mortality risk

Our findings demonstrate that delta-age, defined as the difference between the biological age predicted by the clock and the chronological age, is a strong predictor of all-cause mortality. Positive delta-age values were associated with increased mortality risk, while negative values were predictive of reduced risk. These associations remained robust whether we analyzed delta-age as a continuous or categorical variable. These results align with previous reports of biological clocks capturing aging-related functional decline [6], [17], and they further support the utility of transcriptomic clocks as meaningful biomarkers of aging. The effect sizes observed here (HR = 1.108 per unit delta-age; HR = 1.561 for those >3 years older) underscore the relevance of transcriptome-derived age acceleration not only as a reflection of biological aging but also as a clinically informative metric for mortality risk stratification, assessment of individual health status, and monitoring intervention effects.

### Limitations and Future Directions

#### Blood cell type composition changes

The composition of blood cell types changes with age, thereby influencing gene expression levels in whole blood [66]. For instance, a single-cell study in centenarians observed a decrease in naive CD4 T-cells, memory CD4 T-cells, and naive B-cells, alongside an increase in CD8 T-cells, natural killer cells, and monocytes proportions among the extreme old individuals compared to younger age groups [9].

In this study, we did not incorporate blood cell type information, as our primary objective was to identify easily accessible aging markers in blood, the rationale being that even if a marker’s significance is induced by changes in blood cell type composition, it remains a valid aging marker. Current work is ongoing to further investigate the association between cell type composition with age.

#### Analysis of longitudinal changes

All findings discussed in this study are the results of cross-sectional analyses. A previous longitudinal multi-omics study identified diverse aging patterns among individuals, highlighting that longitudinal changes within individuals can diverge from the cross-sectional trends observed in molecular features [67]. Currently, LLFS is actively collecting and profiling blood transcriptome data at various time points. At the time this manuscript is being prepared, LLFS has collected transcriptomic data from over 1,100 participants across at least two time points, with time intervals ranging from 5 to 17 years. Among them, more than 300 participants have measurements at three time points, providing a valuable dataset for evaluating individual longitudinal changes. As a subsequent step, we will explore longitudinal changes in transcriptomic profiles across participants of different ages, which will offer deeper insight into the dynamics of molecular features over time.

#### Integration of multi-omics, single-cell, and tissue-specific analysis

This study focused exclusively on whole blood transcriptomic analysis. However, aging and mortality risk are complex processes that involve changes across multiple molecular and tissue-specific levels. Future research that integrates multi-omics datasets may yield a more comprehensive understanding of these processes. Moreover, while whole blood offers advantages such as minimally invasive sample collection and relevance in systemic biological studies, it is a heterogeneous tissue and lacks cellular specificity. Analyzing specific tissue types, and further diving into the single-cell space could provide a more detailed and systematic view of the aging process.

## Grant Funding

This research was supported by NIH-NIA U19 AG063893 and UH2/UH3AG064704 (ILO).

## Competing Interest Declaration

The authors declare no competing interests.

## Ethics Declaration

This study was conducted in accordance with the relevant local and international ethical guidelines. All enrolled participants (or their legal guardian or representative, when applicable) provided informed consent. All study personnel and data analysts have completed and maintained current human-subjects research training under IRB oversight. All participant data used, including phenotypical data, genetic data, and RNA-seq data, were de-identified and are stored on secure servers with restricted access.

## Author Contribution Declaration

Analysis, methods and manuscript writing: Mengze Li.

Analysis, methods and manuscript editing: Zeyuan Song, Eric Reed, Tanya T. Karagiannis, Stacy Andersen.

RNA-seq data acquiring and processing: Michael Brent, Chase Mateusiak, Sandeep Acharya, Wooseok . J. Jung, Jeffrey R. O’Connell, Sofiya Milman.

Participant recruiting, administration, phenotype data collection and processing: Shu Liao, Mary K. Wojczynski, Mary F. Feitosa, Konstantin Arbeev, May E. Montasser, Albert Tai, Roland J. Thorpe.

Analysis guidance, ideation, manuscript editing, and funding acquiring: Stefano Monti, Paola Sebastiani, Thomas T. Perls.

---

**Supplement Figure 1.**
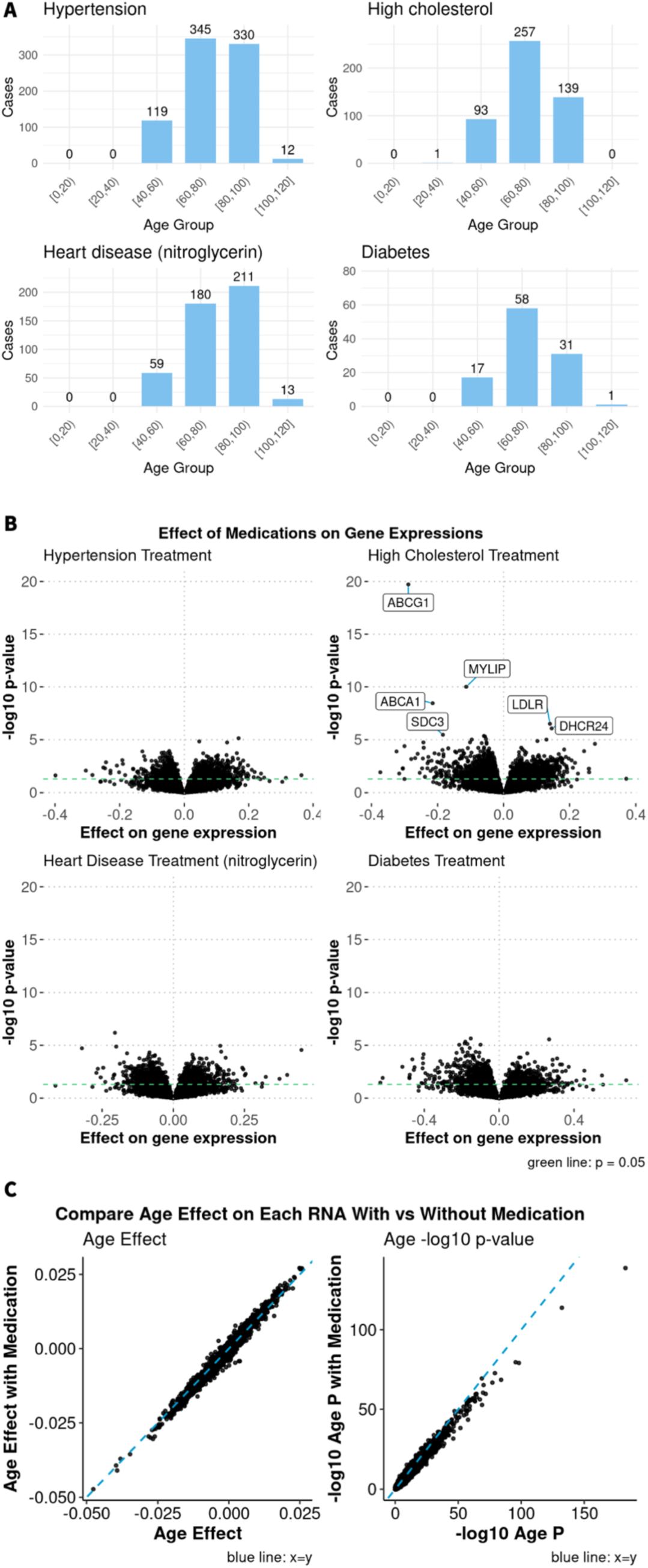
Medication has minimal changes in association between gene expression level and age. A. Bar plots showing the number of participants taking each medication in each age groups. B. Volcano plots showing effect size estimates versus p-values for gene expression associations with four medications, analyzed using the same model as the age association test. Genes significantly associated with medication (FDR q ≤ 0.01) are labeled. The green horizontal dashed line indicates p = 0.05. C. Scatter plots comparing the estimates and the negative log 10 nominal p-values of the analyses with and without medication added to the model. The systematic decrease in p-value significance when including medications is explained by the reduced sample size since only a fraction of the participants have medication information available. The blue dash line represents the identity line x=y.

**Supplement Figure 2.**
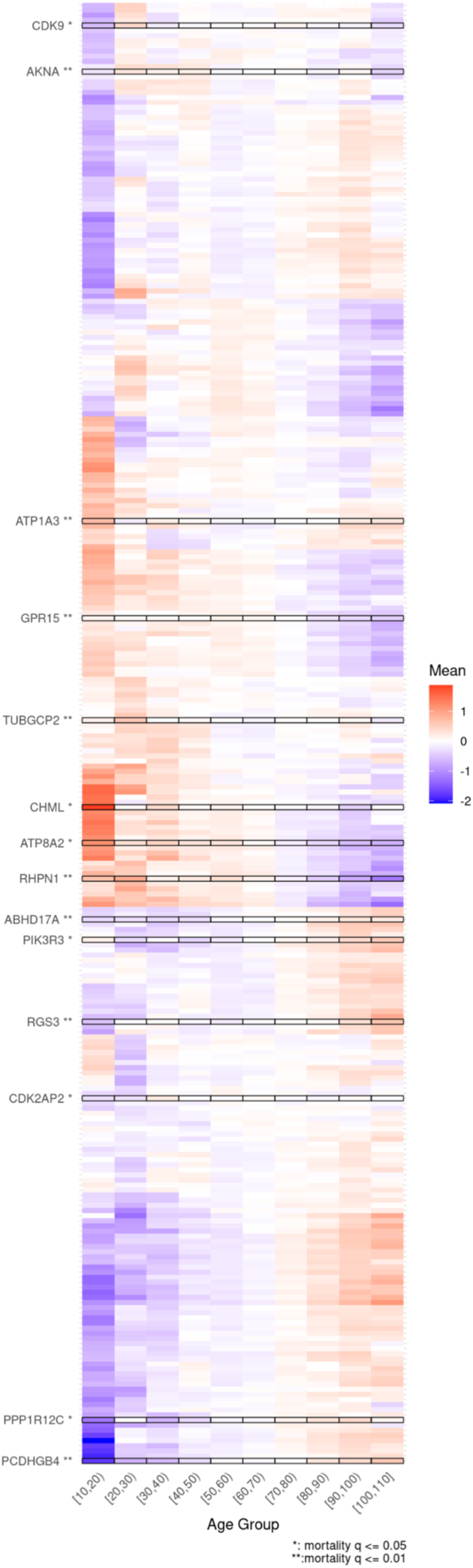
Heatmap of the expression level of the markers exclusively in LLFS and ILO. Heatmap showing the normalized expression levels of 286 gene markers exclusively associated with age in LLFS and ILO cohorts, across different age groups. Genes significantly associated with mortality risk (q ≤ 0.05 and q ≤ 0.01) are highlighted. Genes are clustered using hierarchical clustering based on their expression patterns.

Supplement Table 1. Comprehensive table of all the mRNA associations with age in LLFS and ILO, and mortality risk in LLFS. age_sig are defined by if the gene marker has a q value (FDR corrected) ≤ 0.01. age_dir refers to if the expression of the gene marker increase (Inc) or decrease (Dec) with age.

Supplement Table 2. Significantly enriched gene sets and pathways (q ≤ 0.05) identified from gene markers associated with age (Age_up and Age_dn) and mortality risk (HR > 1 and HR < 1). Enrichment was assessed using a Kolmogorov–Smirnov (KS) test with genes ranked by Z score. label: gene set or pathway names. gene set: total number of genes in this gene set. overlap: the number of genes overlapped between the gene marker list and the gene set. pval and fdr: p-value and q-value (FDR correction) of the enrichment significance. score: enrichment score calculated from fgsea package. hits: the overlapped gene symbols between the gene set and the analyzed data set.

## Notes

### Competing Interest Statement

The authors have declared no competing interest.

### Summary of Updates

Following peer review of this article, we received and addressed the reviewers' comments. We have updated the discussion section to further address the clinical translatability of the identified markers.

